## Supplementary material for "Distinct ventral tegmental area neuronal ensembles are indispensable for reward-driven approach and stress-driven avoidance behaviors": SupplFigs

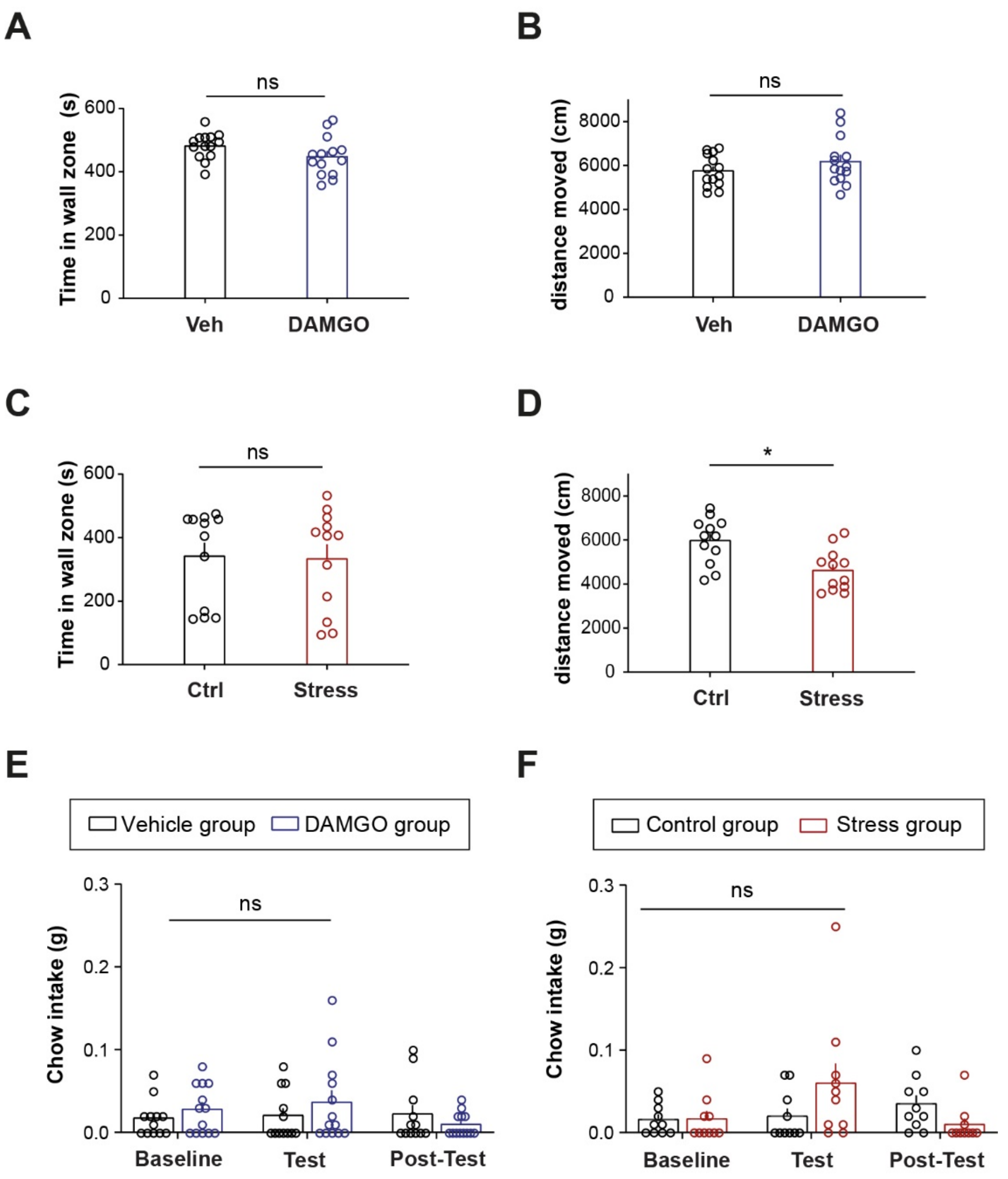


Supplementary Figure 1

**SUPPLEMENTARY FIGURE 1 | (A)** Bar graph showing the average amount of time spent in wall zone during the open field test after vehicle or DAMGO injection (N = 14 per group, unpaired t test, t(26) = 1.699, p = 0.1012). **(B)** Bar graph showing the average total distance moved during the open field test after vehicle or DAMGO injection (N = 14 per group, unpaired t test, t(26) = 1.21, p = 0.2371). **(C)** Bar graph showing the average amount of time spent in wall zone during the open field test after novel cage mate or acute stress exposure (N = 12 per group, unpaired t test, t(22) = 0.1433, p = 0.8874). **(D)** Bar graph showing the average total distance moved during the open field test after novel cage mate or social stress exposure (N = 12 per group, unpaired t test, t(22) = 3.318, p = 0.0031). **(E)** Bar graph showing the average amount of chow consumed during an 1-hour binge session after vehicle or DAMGO administration on test day ($N_{veh}$ = 12, $N_{DAMGO}$ = 13, Two-way RM ANOVA, Day x Ligand interaction F(2,46) = 1.604, p = 0.2122). **(F)** Bar graph showing the average amount of chow consumed during an 1-hour binge session after novel cage mate or acute stress exposure (N = 10 per group, Two-way RM ANOVA, Day x Group interaction, F(2,36) = 4.738, p = 0.0149, no significant post-hoc test).

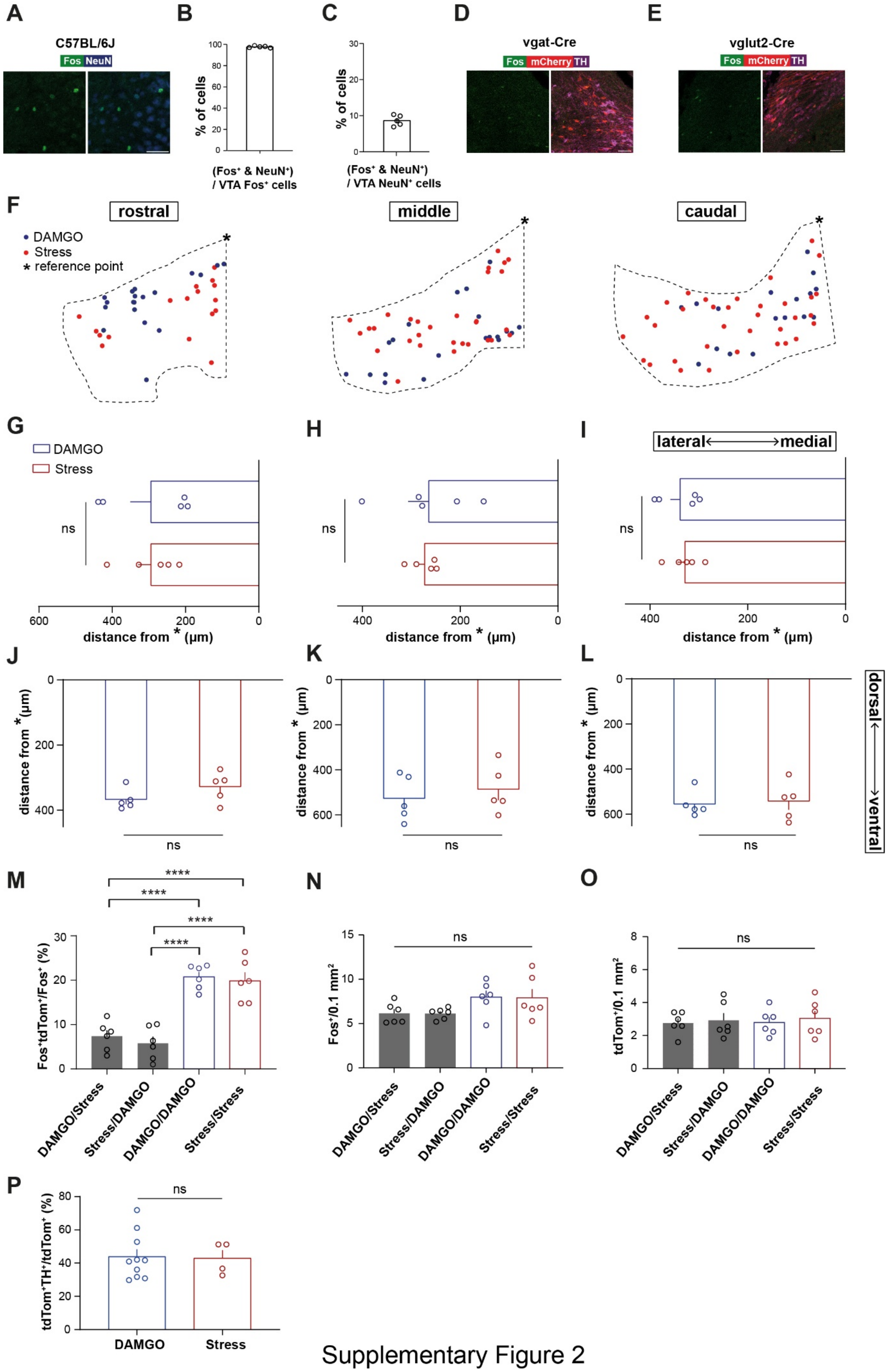


Supplementary Figure 2

**SUPPLEMENTARY FIGURE 2 | (A)** Representative example of Fos/NeuN co-staining in the VTA. Scale bar, 50 microns. **(B)** Bar graph showing the percentage of double $Fos^+NeuN^+$/ $Fos^+$ (N = 5 mice). **(C)** Bar graph showing the percentage of double $Fos^+NeuN^+/NeuN^+$ (N = 5 mice). **(D)** Representative example of triple Fos, mCherry, and TH labeling in the VTA. mCherry is expressed in $VGAT^+$ cells. Scale bar, 50 microns. **(E)** Representative example of triple Fos, mCherry, and TH labeling in the VTA. mCherry is expressed in $Vglut2^+$ cells. Scale bar, 50 microns. **(F)** Representative examples of VTA topography of $Fos^+$ cells after DAMGO (blue) or social stress (red) across rostral, middle and caudal VTA. **(G)** Bar graph showing the average mediolateral distance of $Fos^+$ neurons from a reference point after DAMGO or stress in the rostral VTA (N = 5 per group, unpaired t-test, $t(8) = 0.00172$, $p = 0.9987$). **(H)** Bar graph showing the average mediolateral distance of $Fos^+$ neurons from a reference point after DAMGO or stress in the middle VTA (N = 5 per group, unpaired t-test, $t(8) = 0.1902$, $p = 0.8539$). **(I)** Bar graph showing the average mediolateral distance of $Fos^+$ neurons from a reference point after DAMGO or stress in the caudal VTA (N = 5 per group, unpaired t-test, $t(8) = 0.3951$, $p = 0.7031$). **(J)** Bar graph showing the average dorsoventral distance of $Fos^+$ neurons from a reference point after DAMGO or stress in the rostral VTA (N = 5 per group, unpaired t-test, $t(8) = 1.57$, $p = 0.1550$). **(K)** Bar graph showing the average dorsoventral distance of $Fos^+$ neurons from a reference point after DAMGO or stress in the middle VTA (N = 5 per group, unpaired t-test, $t(8) = 0.6194$, $p = 0.5529$). **(L)** Bar graph showing the average dorsoventral distance of $Fos^+$ neurons from a reference point after DAMGO or stress in the caudal VTA (N = 5 per group, unpaired t-test, $t(8) = 0.2889$, $p = 0.78$). **(M)** Bar graph showing the percentage of double $tdTom^+$ and $Fos^+$ / $Fos^+$ cells (N = 6 per group, One-way ANOVA, $F(3,20) = 28$, $p < 0.0001$, Tukey's post-hoc comparisons: DAMGO/stress vs DAMGO/DAMGO, $q(20) = 8.885$, $p < 0.0001$, DAMGO/stress vs stress/stress, $q(20) = 8.294$, $p < 0.0001$, stress/DAMGO vs DAMGO/DAMGO, $q(20) = 9.955$, $p < 0.0001$, stress/DAMGO vs stress/stress, $q(20) = 9.364$, $p<0.0001$). **(N)** Bar graph showing the average number of $Fos^+$ / 0.1 $mm^2$ (N = 6 per group, One-way ANOVA, $F(3,20) = 2.462$, $p = 0.0922$). **(O)** Bar graph showing the average number of $tdTom^+$ / 0.1 $mm^2$ (N = 6 per group, One-way ANOVA, $F(3,20) = 0.1122$, $p = 0.9519$). **(P)** Bar graph showing the percentage of double $tdTom^+TH^+/tdTom^+$ for $VTA_{DAMGO}$ and $VTA_{STRESS}$ ($N_{DAMGO} = 10$, $N_{STRESS} = 4$, unpaired t test, $t(12) = 0.1231$, $p = 0.9041$).

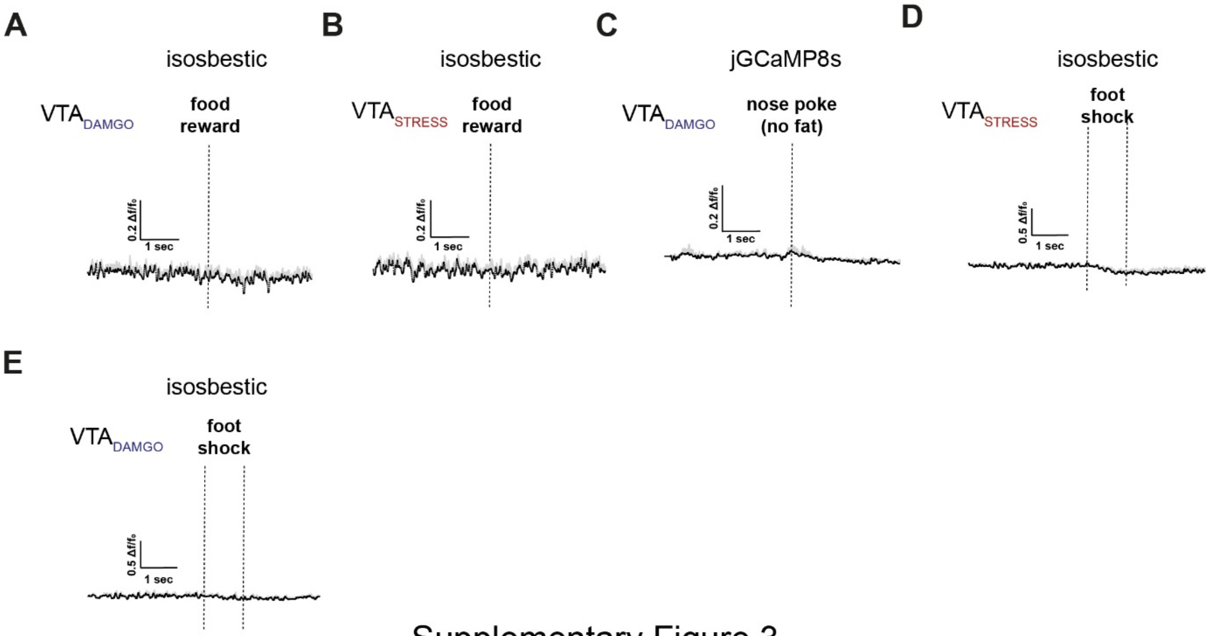


Supplementary Figure 3

**SUPPLEMENTARY FIGURE 3 | (A)** Line plot showing the average $\Delta f/F_0$ of isosbestic control for $VTA_{DAMGO}$ ensemble during food reward. **(B)** Line plot showing the average $\Delta f/F_0$ of isosbestic control for $VTA_{STRESS}$ ensemble during food reward. **(C)** Line plot showing the average $\Delta f/F_0$ jGCaMP8s signal from $VTA_{DAMGO}$ time-locked to nose-poking in an empty port. **(D)** Line plot showing the average $\Delta f/F_0$ of isosbestic control for $VTA_{STRESS}$ ensemble during foot shock. **(E)** Line plot showing the average $\Delta f/F_0$ of isosbestic control for $VTA_{DAMGO}$ ensemble during foot shock.

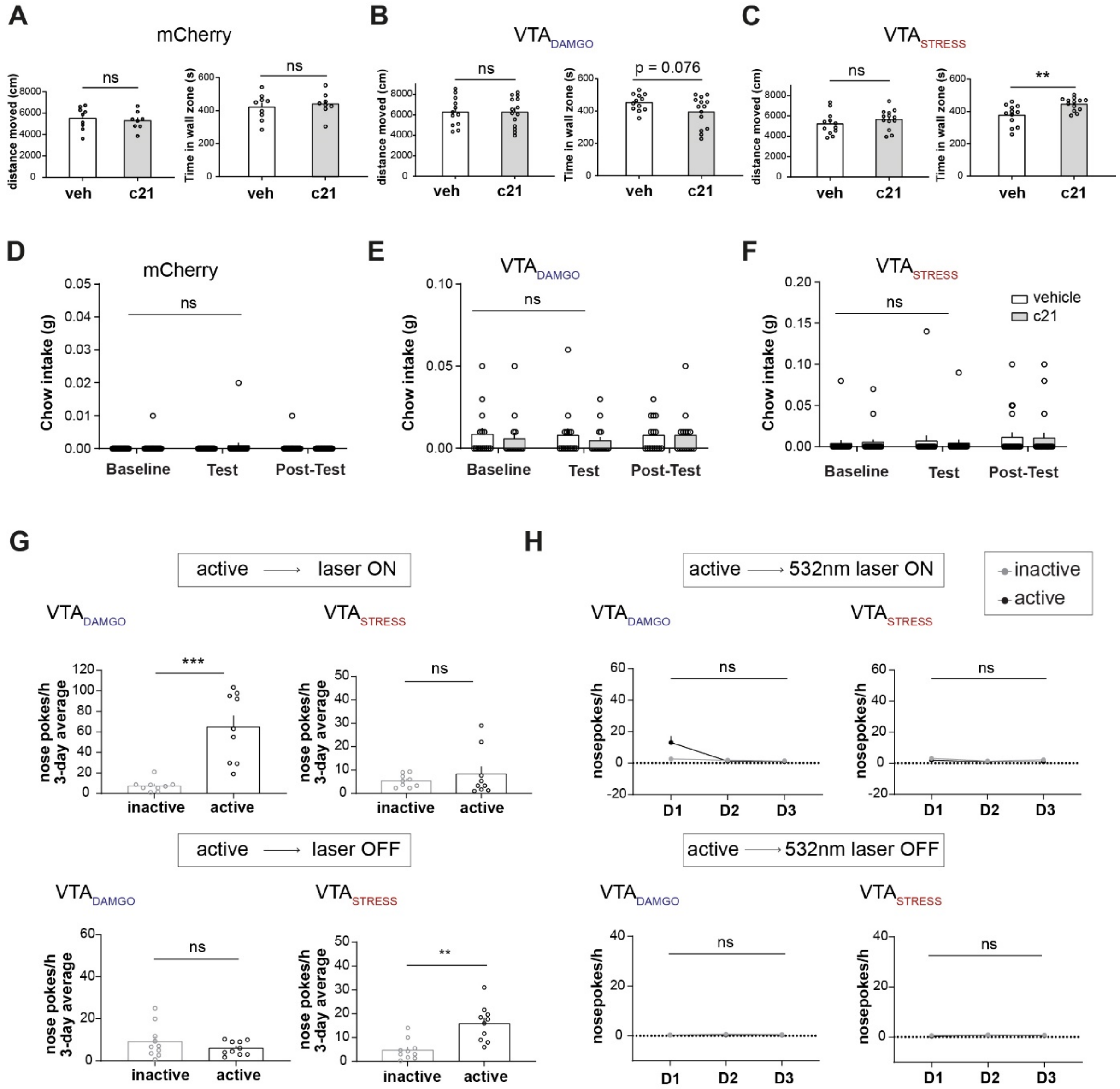


Supplementary Figure 4

**SUPPLEMENTARY FIGURE 4 | (A)** Bar graph showing the average distance moved (left) and time spent in wall zone (right) in the open field test for mCherry control mice after vehicle or c21 in injection ($N_{veh}$ = 9, $N_{c21}$ = 8, distance moved: unpaired t test, t(15) = 0.4842, p = 0.6352, time in wall zone: two unpaired t test, t(15) = 0.4876 , p = 0.6329). **(B)** Bar graph showing the average distance moved (left) and time spent in wall zone (right) in the open field test for the $VTA_{DAMGO}$ group after vehicle or c21 in injection ($N_{veh}$ = 12, $N_{c21}$ = 14, distance moved: unpaired t test, t(24) = 0.0003532, p = 0.09997, time in wall zone: unpaired t test, t(24) = 1.854, p = 0.0761). **(C)** Bar graph showing the average distance moved (left) and time spent in wall zone (right) in the open field test for the $VTA_{STRESS}$ group after vehicle or c21 (1 mg/kg) in injection ($N_{veh}$ = 12, $N_{c21}$ = 13, distance moved: unpaired t test, t(23) = 1.021 ,p = 0.3181, time in wall zone: unpaired t test, t(23) = 3.206 , p = 0.0039). **(D)** Bar graph showing the average chow intake during baseline, test, and post-test days for the mCherry control group. Mice received vehicle or c21 (1 mg/kg) on test day in a counterbalanced manner (N = 22, Two-way RM ANOVA, Day x Ligand interaction, F(2,84) = 0.2749, p = 0.7603). **(E)** Bar graph showing the average chow intake during baseline, test, and post-test days for the $VTA_{DAMGO}$ group. Mice received vehicle or c21 (1 mg/kg) on test day in a counterbalanced manner (N = 15, Two-way RM ANOVA, Day x Ligand interaction, F(2,56) = 0.1318, p = 0.8768). **(F)** Bar graph showing the average chow intake during baseline, test, and post-test days for the $VTA_{STRESS}$ group. Mice received vehicle or c21 (1 mg/kg) on test day in a counterbalanced manner (N = 21, Two-way RM ANOVA, Day x Ligand interaction, F(2,80) = 0.1651, p = 0.8481). **(G)** Top: Bar graphs showing the 3-day average number of nose-pokes to activate a 473nm laser for $VTA_{DAMGO}$ and $VTA_{STRESS}$ groups ($VTA_{DAMGO}$ : N = 9, paired t test, t(8) = 5.622, p = 0.0005 | $VTA_{STRESS}$: N = 10, paired t test, t(9) = 1.446, p = 0.1821). Bottom: Bar graphs showing the 3-day average number of nose-pokes to deactivate a 473nm laser for $VTA_{DAMGO}$ and $VTA_{STRESS}$ groups ($VTA_{DAMGO}$ : N = 9, paired t test, t(8) = 0.8623, p = 0.4136 | $VTA_{STRESS}$: N = 10, paired t test, t(9) = 4.104, p = 0.0027). **(H)** Top: Line plot showing the average number of nose pokes per hour to activate a 532 nm laser for $VTA_{DAMGO}$ and $VTA_{STRESS}$ on D1-D3 ($N_{DAMGO}$ = 9, Two-way RM ANOVA, Day x Nose-poke interaction, F(2,32) = 6.434, p = 0.0045, no significant post-hoc | $N_{STRESS}$ = 10, Two-way RM ANOVA, Day x Nose-poke interaction, F(2,36) = 0.2363, p = 0.7908). Bottom: Line plot showing the average number of nose pokes per hour to deactivate a 532 nm laser for $VTA_{DAMGO}$ and $VTA_{STRESS}$ on D1-D3 ($N_{DAMGO}$ = 9, Two-way RM ANOVA, Day x Nose-poke interaction, $F(2,32) = 0.1231$, $p = 0.8846$ | $N_{STRESS} = 10$, Two-way RM ANOVA, $F(2,36) = 0.1667$, $p = 0.8471$).

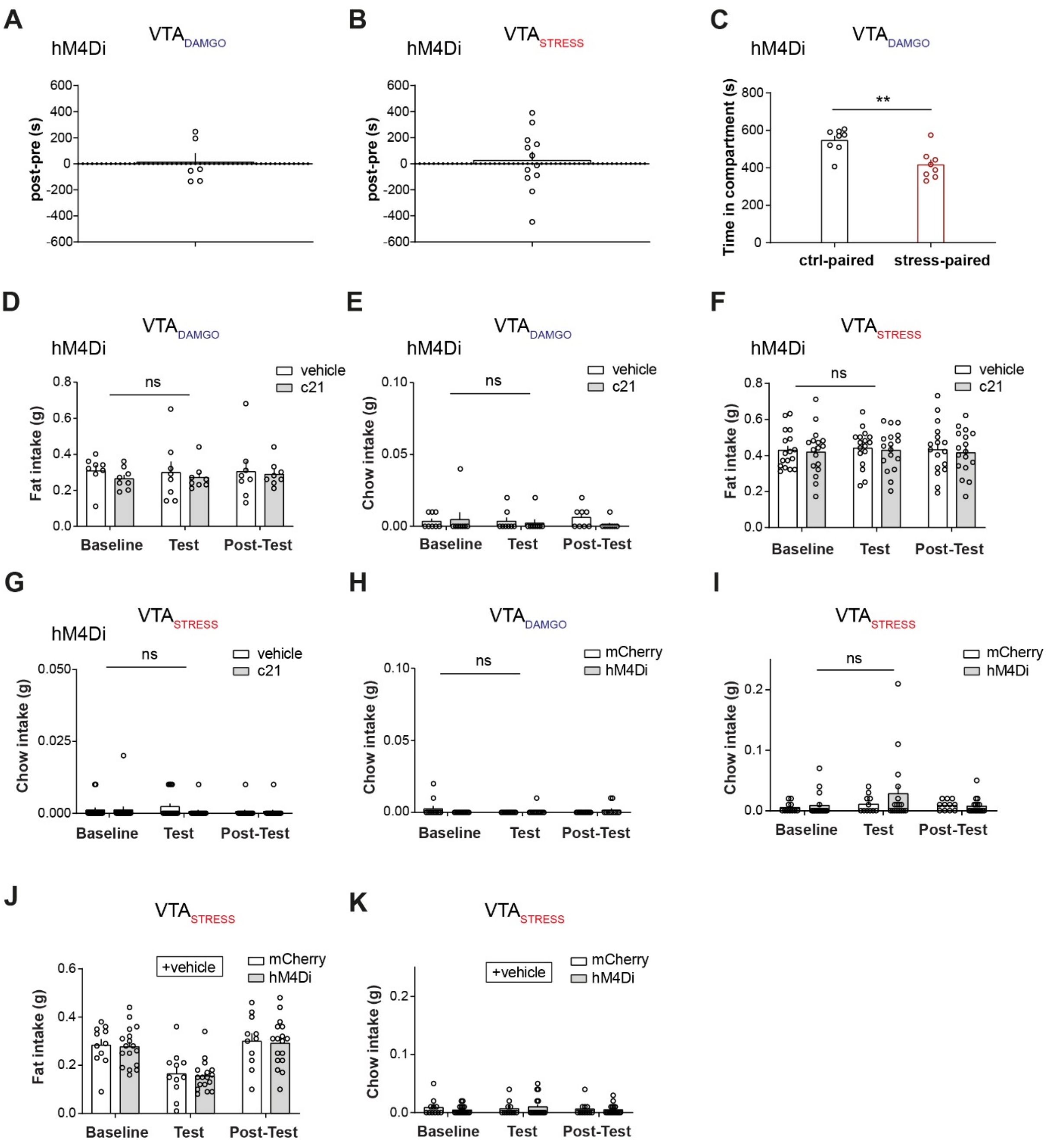


Supplementary Figure 5

**SUPPLEMENTARY FIGURE 5 | (A)** Bar graph showing the average preference score for $VTA_{DAMGO}$ hM4Di-expressing mice (N = 6, paired t test, t(5) = 0.1963, p = 0.8521). **(B)** Bar graph showing the average preference score for $VTA_{STRESS}$ hM4Di-expressing mice (N = 12, paired t test, t(11) = 0.3854, p = 0.7073). **(C)** Bar graph showing the time spent in control-paired or stress-paired compartments in $VTA_{DAMGO}$ hM4Di-expressing animals (N = 8, paired t test, t(14) = 3.599, p = 0.0029). Showing that inhibition of $VTA_{DAMGO}$ ensemble does not affect stress-driven CPA. **(D)** Bar graph showing the amount of fat consumed (g) for $VTA_{DAMGO}$ hM4Di-expressing animals during baseline, test, and post-test days. During test-day mice received vehicle or c21 (2 mg/kg) in a counter-balanced manner before starting the feeding session (N = 8, Two-way RM ANOVA, Day x Ligand interaction, F(2,28) = 0.1662, p = 0.8477). **(E)** Bar graph showing the amount of chow consumed (g) for $VTA_{DAMGO}$ hM4Di-expressing animals during baseline, test, and post-test days. During test-day mice received vehicle or c21 (2 mg/kg) in a counter-balanced manner before starting the feeding session (N = 8, Two-way RM ANOVA, Day x Ligand interaction, F(2,28) = 0.6488, p = 0.5304). **(F)** Bar graph showing the amount of fat consumed (g) for $VTA_{STRESS}$ hM4Di-expressing animals during baseline, test, and post-test days. During test-day mice received vehicle or c21 (2 mg/kg) in a counter-balanced manner before starting the feeding session (N = 17, Two-way RM ANOVA, Day x Group interaction, F(2,64) = 0.008964 , p = 0.9911). **(G)** Bar graph showing the amount of chow consumed (g) for $VTA_{STRESS}$ hM4Di-expressing animals during baseline, test, and post-test days. During test-day mice received vehicle or c21 (2 mg/kg) in a counter-balanced manner before starting the feeding session (N = 17, Two-way RM ANOVA, Day x Group interaction, F(2,64) = 0.8889 , p = 0.4161). **(H)** Bar graph showing the amount of chow consumed (g) during baseline, test, and post-test day for $VTA_{DAMGO}$ mCherry- or hM4Di-expressing mice. During test day mice received an injection of c21 (2 mg/kg) followed by an injection of DAMGO (1 mg/kg) before the beginning of the feeding session ($N_{mCherry}$ = 11, $N_{hM4Di}$ = 11, Day x Group interaction, Two-way RM ANOVA, F(2,40) = 3.231, p = 0.05). **(I)** Bar graph showing the amount of chow consumed (g) during baseline, test, and post-test day for $VTA_{STRESS}$ mCherry- or hM4Di-expressing mice. During test day mice received an injection of c21 (2 mg/kg) followed by a 20 sec episode of social stress before the beginning of the feeding session ($N_{mCherry}$ = 11, $N_{hM4Di}$ = 17, Two-way RM ANOVA, Day x Group interaction, F(2,52) = 0.9741, p = 0.3843). **(J)** Bar graph showing the amount of fat consumed (g) during baseline, test, and post-test days for $VTA_{STRESS}$ mCherry- and hM4Diexpressing mice. During test day mice received a vehicle injection followed by a 20 s episode of social stress prior to the beginning of the feeding session ($N_{mCherry}$ = 11, $N_{hM4Di}$ = 17, Two-way RM ANOVA, Day main effect, $F(1.818,47.27) = 36.35$, $p < 0.0001$). **(K)** Bar graph showing the amount of chow consumed (g) during baseline, test, and post-test days for $VTA_{STRESS}$ mCherry- and hM4Di-expressing mice. During test day mice received a vehicle injection followed by a 20 s episode of social stress prior to the beginning of the feeding session ($N_{mCherry}$ = 11, $N_{hM4Di}$ = 17, Two-way RM ANOVA, Day x Group interaction, $F(2,52) = 0.5835$, $p = 0.5615$).
